# Deconstructing Procedural Memory: Different Learning Trajectories and Consolidation of Sequence and Statistical Learning

**DOI:** 10.1101/216374

**Authors:** Peter Simor, Zsofia Zavecz, Kata Horváth, Noémi Éltető, Csenge Török, Orsolya Pesthy, Ferenc Gombos, Karolina Janacsek, Dezso Nemeth

## Abstract

Procedural learning is a fundamental cognitive function that facilitates efficient processing of and automatic responses to complex environmental stimuli. Here, we examined training-dependent and off-line changes of two sub-processes of procedural learning: namely, sequence learning and statistical learning. Whereas sequence learning requires the acquisition of order-based relationships between the elements of a sequence, statistical learning is based on the acquisition of probabilistic associations between elements. Seventy-eight healthy young adults (58 females and 20 males) completed the modified version of the Alternating Serial Reaction Time task that was designed to measure Sequence and Statistical Learning simultaneously. After training, participants were randomly assigned to one of three conditions: active wakefulness, quiet rest, or daytime sleep. We examined off-line changes in Sequence and Statistical Learning as well as further improvements after extended practice. Performance in Sequence Learning increased during training, while Statistical Learning plateaued relatively rapidly. After the off-line period, both the acquired sequence and statistical knowledge was preserved, irrespective of the vigilance state (awake, quiet rest or sleep). Sequence Learning further improved during extended practice, while Statistical Learning did not. Moreover, within the sleep group, cortical oscillations and sleep spindle parameters showed differential associations with Sequence and Statistical Learning. Our findings can contribute to a deeper understanding of the dynamic changes of multiple parallel learning and consolidation processes that occur during procedural memory formation.

## Introduction

Procedural learning, the development of perceptual and motor skills through extensive practice is a crucial ability that facilitates efficient processing of and automatic responses to complex environmental stimuli. Procedural learning is evidenced by enhanced performance as well as functional changes in the neural network underlying behavior (Howard et al., 2004; Fletcher et al., 2005). Learning performance does not only depend on training during acquisition but also on the post-learning period (Karni et al., 1998; Doyon et al., 2009; Durrant et al., 2011). Nevertheless, there are intensive debates questioning whether the acquired memories are stabilized or enhanced during post-learning, off-line periods (Maquet et al., 2000; Krakauer and Shadmehr, 2006; Peigneux et al., 2006; Rickard et al., 2008; Doyon et al., 2009; Nemeth et al., 2010; Pan and Rickard, 2015; Mantua, 2018). Mixed findings emerging in this field suggest that different processes *within* the procedural learning domain may show different trajectories during learning and off-line periods. At least two processes underlying procedural learning can be distinguished: sequence learning and statistical learning (Nemeth et al., 2013; Kóbor et al., 2018). Sequence learning refers to the acquisition of a series of (usually 5-12) stimuli that repeatedly occur in the same *order* (with no embedded noise in deterministic sequences, or with some embedded noise in probabilistic sequences). In contrast, statistical learning refers to the acquisition of shorter-range relationships among stimuli that is primarily based on *frequency* information (i.e., differentiating between more frequent and less frequent runs (e.g., pairs, triplets, etc.) of stimuli. Previous research has not directly contrasted the consolidation of these two processes. Here, we show - using a visuomotor probabilistic sequence learning task - that performance in sequence learning compared to statistical learning (acquisition of order vs. frequency information) shows marked practice-dependent improvements before and after off-line periods.

Studies on sequence learning showed enhanced behavioral performance after an off-line period spent asleep compared to an equivalent period spent awake, especially if individuals acquired an explicit, abstract or complex representation of the sequence (Robertson et al., 2004; Spencer et al., 2006; King et al., 2017). On the other hand, learning probabilistic sequences (Song et al., 2007a, Nemeth et al., 2010), in contrast to deterministic ones, does not seem to benefit from post-learning sleep on the behavioral level, while on a neural level, it has been shown that post-learning sleep is involved in the reprocessing and optimization of the acquired probabilistic sequential information (Peigneux et al., 2003). Importantly, in these probabilistic sequence learning studies the behavioral index of learning encompassed the acquisition of both order- and frequency-based information, thus, the consolidation of sequence learning and statistical learning was not examined separately (Song et al., 2007a, 2007b; Nemeth et al., 2010). There are several studies that investigated the long term retention of statistical learning (Kim et al., 2009; Nemeth et al., 2010; Kóbor et al., 2017), and there is limited evidence that statistical learning in the auditory domain benefits from sleep (Durrant et al., 2011, 2013). Nevertheless, the consolidation, and more specifically, the role of sleep in statistical learning within the visuomotor domain remains largely unexplored.

The Alternating Serial Reaction Time (ASRT) task is a unique tool to investigate statistical and sequence learning within the same experiment (Howard and Howard, 1997; Nemeth et al., 2013). In this perceptual-motor four-choice reaction time (RT) task, participants are required to respond to visual stimuli appearing on the screen. In this task, predetermined sequential (termed as pattern) trials alternate with random ones (e.g., 2R4R3R1R, where numbers correspond to the four locations on the screen presented in the same sequential order during the entire task, and the letter R represents randomly chosen locations) that results in some chunks of stimuli being more frequent than others (see Fig. 1) and enables us to measure the acquisition of both order and frequency information. Namely, sequence learning is defined as acquiring order information, in that consecutive elements in the sequence (denoted with numbers in the above example) can be predicted with 100% certainty based on the previous sequence element (i.e., the 2^nd^ order transitional probability for the sequence trials is equal to one), while random trials are unpredictable (random stimuli can occur at any of the four possible locations with the same probability). However, as mentioned above, the alternating stimulus structure also results in some chunks of stimuli (three consecutive trials, called *triplets*) occurring more frequently than others (62.5% vs. 12.5%, respectively). For instance, the triplet 2X4 (where X denotes any location out of the four possible ones) would occur more frequently as its first and third item can originate either from sequential/pattern or random stimuli. In contrast, the triplet 2X1 would occur less frequently as this combination can originate only from random stimuli (for more details see Fig 1 and the Methods section). Statistical learning is defined as acquiring this frequency information (which also represents a 2^nd^ order regularity, where the transitional probability is less than one; for more detailed explanation see (Kóbor et al., 2018). To disentangle sequence and statistical learning in the ASRT task, sequence learning is assessed by contrasting sequential/pattern and random stimuli, while controlling for frequency information (i.e., analyzing only high-frequency trials). In contrast, statistical learning is assessed by contrasting high- vs. low-frequency trials while controlling for order information (i.e., analyzing only the random trials) (Nemeth et al., 2013; Kóbor et al., 2018). The learning trajectories for both sequence and statistical learning can be tracked by how different behavioral indices, such as RT and accuracy, change over the course of the task (Howard et al., 2004; Nemeth et al., 2013). To the best of our knowledge, no study has yet tracked the temporal dynamics of learning sequential structures (order information) as well as statistical probabilities (frequency information) within the same experimental design focusing not only on the learning phase but also on consolidation and on further performance changes in a post-consolidation testing phase.

**Figure 1.**
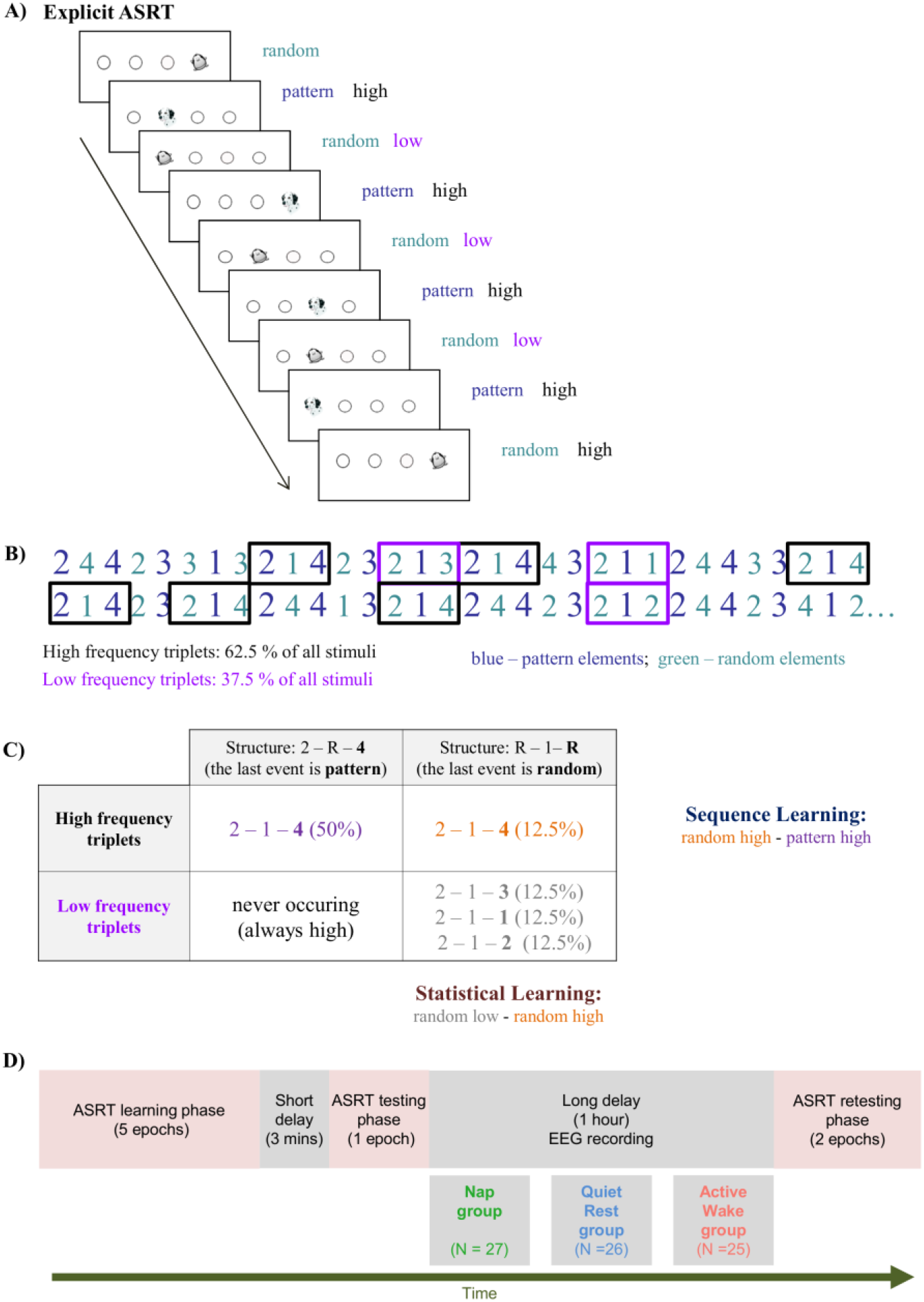
The modified Alternating Serial Reaction Time (ASRT) task. **A)** Pattern and random trials are presented in an alternating fashion. Pattern trials are marked with a picture of a dog, random ones with that of a penguin. Pattern trials always appear in a given location with high probability. Random trials include trials that appear in a given location with high probability and trials that appear in a given location with low probability. **B)** As the ASRT task contains an alternating sequence structure (e.g., 2R4R3R1R, where numbers correspond to the four locations on the screen and the letter R represents randomly chosen locations), some runs of three consecutive elements (called triplets) occur more frequently than others. For subsequent analyses, we determined for each stimulus whether it was the last element of a high-frequency triplet (black frames) or the last element of a low-frequency triplet (purple frames). **C)** We assessed *Statistical Learning* by comparing the responses for those random elements that were the last elements of a high frequency triplet, opposite to those that were the last of a low frequency triplet. In contrast, *Sequence Learning* was quantified as the difference between responses for pattern elements (which were always high frequency triplets) vs. random-high frequency triplet elements. **D)** Study Design. The training phase consisted of five epochs (25 blocks). The testing and retesting phases comprised one and two (that is, 5 and 10 blocks), respectively.

Although sequence learning and statistical learning seem to require different cognitive mechanisms (Nemeth et al., 2013) in everyday learning scenarios, humans might rely simultaneously on both forms of learning. Nevertheless, previous studies investigated the consolidation of these processes in separate task conditions. Therefore, the first aim of our study was to examine the consolidation of sequence learning and statistical learning simultaneously, in the same experimental context. Previous studies suggest that sequence learning may, whereas statistical learning may not benefit from post-learning sleep or more specific oscillatory activity (slow wave activity and spindles); however, these studies applied awake control groups engaged in daytime activities during the off-line periods (King et al., 2017).

As the amount of interference might influence off-line memory processing (Mednick et al., 2011), our second aim was to examine the off-line change of sequence learning and statistical learning after three different post-learning conditions: active wakefulness, quiet rest, and daytime sleep. We hypothesized that sequence learning would be enhanced after sleep and quiet rest (i.e., due to low interference) compared to active wakefulness, whereas off-line change in statistical learning would be independent from the post-learning condition.

Although post-learning sleep seem to facilitate learning capacity in different cognitive domains (Feld and Diekelmann, 2015), several studies indicate that not sleep *per se*, but specific oscillations during sleep facilitate post-sleep improvements in behavioral performance (Rasch and Born, 2013). Among these oscillations, slow waves and sleep spindles emerge as important candidates that reflect processes of memory consolidation and synaptic plasticity (Diekelmann and Born, 2010; Fogel and Smith, 2011; Ulrich, 2016). Slow waves around 1 Hz and especially fast sleep spindles (13-16 Hz) are considered as hallmarks of the reactivation and neocortical redistribution of hippocampus-dependent memories (Diekelmann and Born, 2010). In addition, slow frequency oscillations ranging between 1-8 Hz were linked to the restorative (homeostatic) function of sleep (Achermann et al., 1993; Marzano et al., 2010). In order to examine the associations between cortical oscillations and behavioral performance, we explored the EEG correlates of off-line changes in sequence and statistical learning. We hypothesized that slow frequency oscillations and fast sleep spindles within the sleep group would be positively associated with the post-sleep gains in sequence learning, but not with those of statistical learning.

## Materials and methods

### Participants

Participants (all native Hungarians) were selected from a large pool of undergraduate students from the Eötvös Loránd University in Budapest. The first step of the selection procedure consisted of the completion of an online questionnaire assessing sleep quality and mental health status. Sleep-related questionnaires included the Pittsburgh Sleep Quality Index (PSQI, Buysse et al., 1989; Takács et al., 2016), and Athens Insomnia Scale (AIS, Soldatos et al., 2003; Novák, 2004). Participants that showed poor sleep quality based on previous normative measurements were not included. The Hungarian version of the short (nine item) Beck Depression Inventory (BDI, Rózsa et al., 2001) was used to exclude participants with signs of mild to moderate/severe depression, therefore, participants only with a score less than 10 were included. Respondents reporting current or prior chronic somatic, psychiatric or neurological disorders, or the regular consumption of pills other than contraceptives were also excluded. In addition, individuals reporting the occurrence of any kind of extreme life event (e.g., accident) during the last three months that might have had an impact on their mood, affect and daily rhythms were not included in the study. Only right-handed individuals as verified by the Edinburgh handedness inventory (Oldfield, 1971) were invited to the laboratory. At the first encounter with the assistant, participants were instructed to follow their usual sleep-wake schedules during the week prior to the experiment and to refrain from consuming alcohol and all kinds of stimulants 24 hours before the day of the experiment. Sleep schedules were monitored by sleep agendas, as well as by the adapted version of the Groningen Sleep Quality Scale (Simor et al., 2009) in order to assess individuals’ sleep quality the night before the experiment. The data of participants reporting poor sleep quality the night before the experiment (> 7 points) were not considered in the analyses.

After the above selection procedure, 96 right-handed (28 males, *M*_age_ = 21.66±1.98) participants with normal or corrected-to-normal vision were included in the study. Participants were randomly assigned to one of three groups: an Active Wake, a Quiet Rest, or a Nap group. Individuals unable to fall asleep in the Nap group (*N* = 10) as well as those falling asleep in the awake groups (*N* = 5) were excluded from the final analyses. Furthermore, 3 additional participants were excluded due to the absence of learning in the training session. Therefore, the final behavioral analyses were based on the data of 78 participants (20 males, *M*_age_ = 21.71±1.97), with 25, 26, and 27 participants in the Active Wake, Quiet Rest, and Nap group, respectively (see Table 1). In case of the EEG analyses, the data of 12 participants was excluded due to technical artifacts rendering EEG recordings less reliable. Therefore, physiological analyses were restricted to EEG data with sufficient quality (Active Wake, *N* = 20; Quiet Rest, *N* = 21, Nap, *N* = 25). All participants provided written informed consent before enrollment and received course credits for taking part in the experiment. The study was approved by the research ethics committee of the Eötvös Loránd University, Budapest, Hungary (2015/279). The study was conducted in accordance with the Declaration of Helsinki.

**Table 1.**
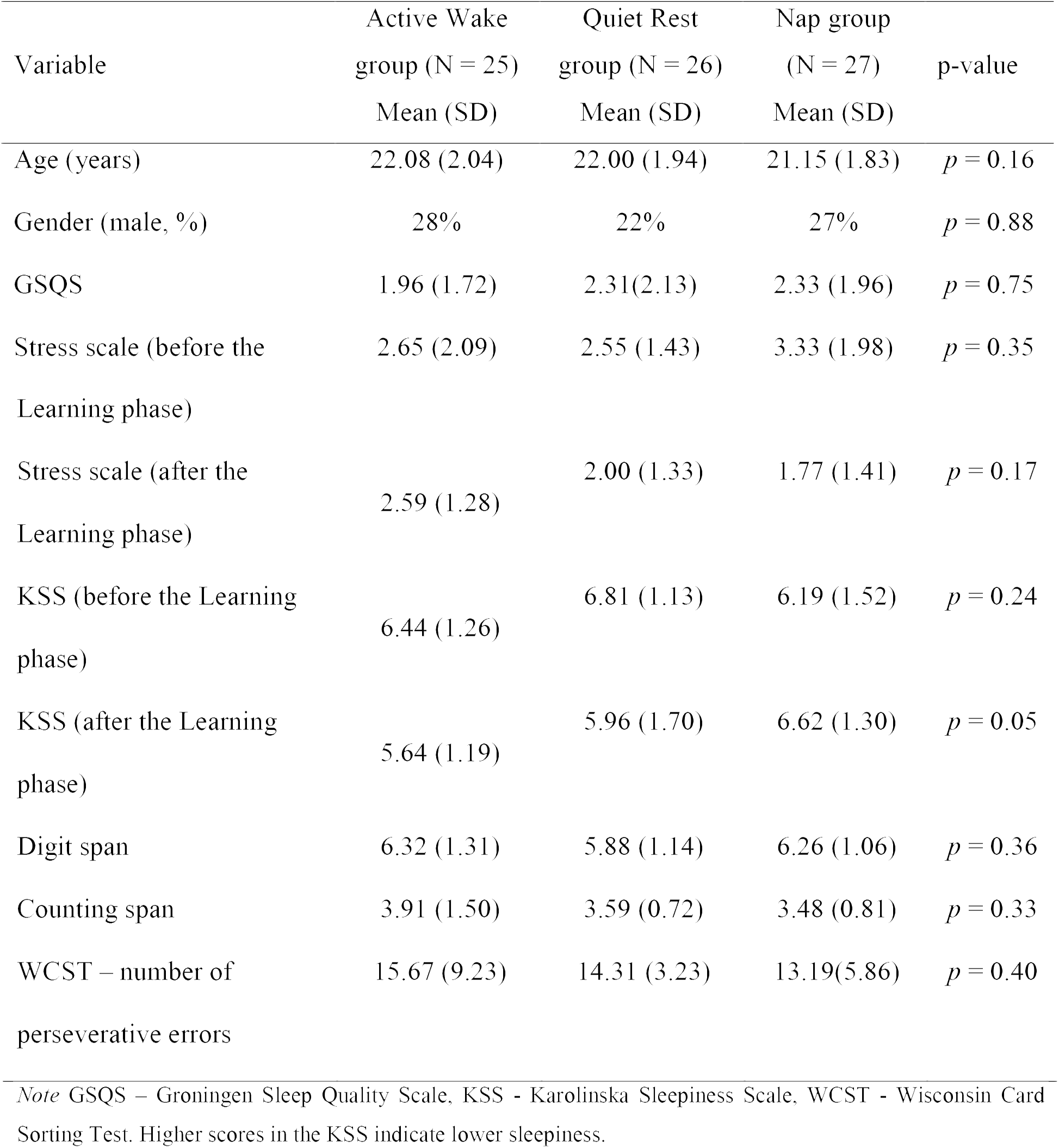
Descriptive characteristics of groups

### Task

Behavioral performance was measured by the explicit version of the Alternating Serial Reaction Time (ASRT) task (Fig. 1, Nemeth et al., 2013). In this task, a stimulus (a dog’s head, or a penguin) appeared in one of four horizontally arranged empty circles on the screen, and participants had to press the corresponding button (of a response box) when it occurred. Participants were instructed to respond as fast and accurate as they could. The task was presented in blocks with 85 stimuli. A block started with five random stimuli for practice purposes, followed by an 8-element alternating sequence that was repeated ten times. The alternating sequence was composed of fixed sequence (pattern) and random elements (e.g., 2-R-4-R-3-R-1-R, where each number represents one of the four circles on the screen and “R” represents a randomly selected circle out of the four possible ones). The response to stimulus interval was set to 120 ms (Song et al., 2007a; Nemeth et al., 2010). In the explicit ASRT task participants are informed about the underlying structure of the sequence, and their attention is drawn to the alternation of sequence and random elements by different visual cues. In our case, a dog always corresponded to sequence elements, and a picture of a penguin indicated random elements (Figure 1A). Participants were informed that penguin targets had randomly chosen locations whereas dog targets always followed a predetermined pattern. They were instructed to find the hidden pattern defined by the dog in order to improve their performance. For each participant, one of the six unique permutations of the four possible ASRT sequence stimuli was selected in a pseudo-random manner, so that the six different sequences were used equally often across participants (Howard and Howard, 1997; Nemeth et al., 2010).

The task consisted of a total of 40 blocks. Participants completed 25 blocks during the *training phase*. As the relatively long training phase can introduce fatigue leading to a general decline in performance measures (e.g., slower reaction times at the end of the training phase that do not reflect the acquired knowledge but the effect of fatigue), a retesting session after a long delay (spent asleep or in wakefulness) can result in a spurious increase in performance because of the release from fatigue. This way, the measure of off-line consolidation is confounded by the effect of fatigue (or more specifically, the release from fatigue) (Pan and Rickard, 2015). In order to control for this factor, the training session was followed by a short (3 minutes long) break in order to minimize the fatigue effect due to massed practice (Rickard et al., 2008; Rieth et al., 2010). After the break, participants were tested on the task for 5 more blocks that constituted the *testing phase*. Subsequently, participants spent an approximately one-hour long off-line period in one of the three conditions (Active Wake, Quiet Rest, and Nap). Finally, they completed a *retesting phase*: 10 more blocks of the same task.

The training phase lasted approximately 30 minutes, the testing phase 5 minutes, and the retesting phase 10 minutes. Awareness of the sequence (pattern elements) was measured after each block. Participants had to type in the regularities they noticed during the task using the same response buttons they used during the ASRT blocks. This method allowed us to determine the duration (in terms of the number of blocks) participants needed to learn the sequence correctly as defined by consistently reporting the same sequence from that point on in the remaining blocks.

### Trial types and learning indices

The alternating sequence of the ASRT task forms a sequence structure in which some of the runs of three successive elements (henceforth referred to as triplets) appear more frequently than others. In the above example, triplets such as 2X4, 4X3, 3X1, and 1X2 (X indicates the middle element of the triplet) occur frequently since the first and the third elements can either be pattern or random stimuli. However, 3X2 and 4X2 occur less frequently since the first and the third elements can only be random stimuli. Figure 1B and 1C illustrate this phenomenon with the triplet 2-1-4 occurring more often than other triplets such as 2-1-3, 2-1-1, and 2-1-2. The former triplet types are labeled as *high-frequency* triplets whereas the latter types are termed as *low-frequency* triplets (see Figure 1C and Nemeth et al., 2013).

The third element of a high-frequency triplet is highly predictable (with 62.5 % probability) from the first element of the triplet. In contrast, in low-frequency triplets the predictability of the third element is much lower (based on a probability of 12.5 %). According to this principle, each stimulus was categorized as either the third element of a high- or a low-frequency triplet. Moreover, trials are differentiated by the cues (dog and penguin) indicating whether the stimulus belongs to the pattern or the random elements. In case of pattern trials, participants can use their explicit knowledge of the sequence to predict the trial, thus we differentiate high-frequency triplets with the last element being a pattern from those triplets in which the last one is a random element. This way, the task consists of three trial types: 1) elements that belong to the explicit sequence and at the same time appear as the last element of a high-frequency triplet are called *pattern* trials; 2) random elements that appear as the last element of a high-frequency triplet are called *random high* trials; and 3) random elements that appear as the last element of a low-frequency triplet are termed *random low* trials (see the example in Figure 1C).

To disentangle the two key learning processes underlying performance on the explicit ASRT task, we differentiate *Sequence Learning* and *Statistical Learning* (Figure 1C). *Sequence Learning* is measured by the difference in reaction times (RT) between random high and pattern elements (the average RT for random high elements minus the average RT for pattern elements). These elements share the same statistical properties (both correspond to the third element of high-frequency triplets), but have different sequence properties (i.e., pattern vs. random elements). Thus, greater Sequence Learning is determined as faster responses to pattern in contrast to random high trials. *Statistical Learning* is assessed by comparing the responses for those random elements that were the last elements of a high-frequency triplet, opposite to those that were the last of a low-frequency triplet (the average RT for random low elements minus the average RT for random high elements). These elements share the same sequence properties (both are random) but differ in statistical properties (i.e., they correspond to the third element of a high or a low-frequency triplet). Hence, faster responses to random high compared to random low trials yields greater Statistical Learning. In sum, Sequence Learning quantifies the advantage (in terms of RT) due to the awareness of the sequential pattern, whereas Statistical Learning captures purely frequency-based learning (Nemeth et al., 2013).

### Procedure

One to two weeks prior the experiment, participants were invited to the laboratory in order to familiarize them with the environment, and to assess their working memory and executive functions based on the Wisconsin Card Sorting Test (PEBL’s Berg Card Sorting Test; Fox et al., 2013) and the Digit Span (Racsmány et al., 2005) and Counting Span (Conway et al., 2005) tasks, respectively. Participants were instructed to complete sleep agendas reporting the schedules, duration and subjective quality of their sleep. On the day of the experiment, participants arrived at the laboratory at 10.00 AM. They completed the GSQS assessing previous nights’ sleep quality. Additionally, their subjective stress levels scored on a 10-point Likert scale (“On a scale from 0-10 how stressed are you feeling now?”), as well as an item of the Hungarian version of the Karolinska Sleepiness Scale (KSS, Akerstedt and Gillberg, 1990) to measure subjective sleepiness were administered. In the Hungarian version of the scale higher scores indicate a more refreshed state, that is, lower sleepiness. Subsequently, EEG caps with 64 electrodes were fitted by two assistants. Testing started at 11.30 AM and took place in a quiet room equipped with a large computer screen, a response box and EEG recording device. After listening to the instructions, participants had the opportunity to practice the task in order to get familiar with the stimuli and the response box; however, all stimuli appeared in a random fashion during the practice session.

This was followed by the explicit ASRT task composed of the *training phase, testing phase, off-line period*, and *retesting phase* (Figure 1D). In the ASRT task, short breaks were introduced between blocks in the following way: First, at the end of each block, participants were instructed to report the sequence they encountered in that block (which took approximately 6 seconds on average). Second, they received feedback for their accuracy and RT performance on pattern trials (fixed 3 seconds). Third, participants were notified (for a fixed 1 second) that the next block can be started by pressing a response button when they are ready; on average, participants continued the next block after approximately 4 seconds. These breaks were somewhat longer for every fifth blocks (i.e., Block 5, 10, 15, etc.), where participants were instructed to continue the next block after EEG data were saved by the experimenter (which took approximately 20 seconds on average). Thus, altogether, for the majority of blocks the between-block break was ~ 14 seconds, and for every fifth block it was ~ 29 seconds. Additionally, a 3-min long break was inserted between the learning and the testing phases during which the fitting of the EEG caps were monitored and impedances were reset under 10 kΩ.

The off-line period extended from 12.30 to 13.30. Participants assigned to the Active Wake group were instructed to watch an approximately one-hour long documentary. (They were allowed to select from documentaries of different topics such as natural sciences, nature or history). Participants of the Quiet Rest group were asked to sit quietly with eyes closed in a comfortable chair. They were instructed by the assistant to open their eyes for 1 minute, every 5 minutes or in case the EEG recording showed any sign of sleep onset (slow eye movements, attenuation of alpha waves and presence of theta oscillations). Participants in the Nap group had the opportunity to spend a daytime nap in the laboratory. The off-line period took place (in all groups) at the same room in which learning, testing and retesting occurred, and was monitored by EEG. Before the retesting phase, participants were asked to complete again the KSS and the scale assessing the level of stress.

### EEG recording

EEG activity was measured by using a 64-channel recording system (BrainAmp amplifier and BrainVision Recorder software, BrainProducts GmbH, Gilching, Germany). The Ag/AgCl sintered ring electrodes were mounted in an electrode cap (EasyCap GmbH, Herrsching, Germany) on the scalp according to the 10% equidistant system. During acquisition, electrodes were referenced to a scalp electrode placed between Fz and Cz electrodes. Horizontal and vertical eye movements were monitored by EOG channels. Three EMG electrodes to record muscle activity, and one ECG electrode to record cardiac activity were placed on the chin and the chest, respectively. All electrode contact impedances were kept below 10 kΩ. EEG data was recorded with a sampling rate of 500 Hz, band pass filtered between (0.3 and 70 Hz).

In order to remove muscle and eye movement related artifact from the awake EEG data (Active Wake and Quiet Rest groups), EEG preprocessing was performed using the Fully Automated Statistical Thresholding for EEG artifact Rejection (FASTER) toolbox (http://sourceforge.net/projects/faster, Nolan et al., 2010) implemented in EEGLAB (Delorme and Makeig, 2004) under Matlab (The Mathworks). The data was first re-referenced to the Fz electrode, notch filtered at 50 Hz, and band-pass filtered between 0.5 – 45 Hz. Using a predefined z-score threshold of ±3 for each parameter, artifacts were detected and corrected regarding single channels, epochs, and independent components (based on the infomax algorithm Bell and Sejnowski, 1995). This way, data was cleared from eye-movement, muscle and heartbeat artifacts. The data was then re-referenced to the average of the mastoid electrodes (M1 and M2). Remaining epochs containing artifacts were removed after visual inspection on a 4-seconds long basis. In case of the sleep recordings (Nap group), data was re-referenced to the average of the mastoid electrodes, and sleep stages as well as conventional parameters of sleep macrostructure were scored according to standardized criteria (Berry et al., 2012) by two experienced sleep researchers. Periods of NREM sleep (Stage 2 and SWS) were considered for subsequent analyses. Epochs containing artifacts were visually inspected and removed on a 4-seconds basis. Wrong channels (N = 6 in the dataset of the Nap group) were replaced by the average of the neighboring channels.

Spectral power and sleep spindle analyses of artifact-free segments were performed by a custom made software tool for EEG analysis (FerciosEEGPlus, © Ferenc Gombos 2008-2017). Overlapping (50%), artifact-free, four-second-epochs of all EEG derivations were Hanning-tapered and Fourier transformed by using the FFT (Fast Fourier Transformation) algorithm in order to calculate the average power spectral densities. The analyzed frequencies spanned between 0.75-31 Hz in the Nap group, and between 1.5 – 25 Hz in the awake groups. Low frequencies (0.75-1.5 Hz) were not considered in the awake conditions due to the negligible and unreliable contribution of measurable cortical activity at this frequency range during wakefulness. In addition, frequencies above 25 Hz were unreliable in the awake data due to technical and movement-related artifacts. We summed up frequency bins to generate five frequency bands for the wake groups: delta (1.5-4 Hz), theta (4.25-8), alpha (8.25-13), sigma (13.25-16), and beta (16.25-25 Hz) frequency bands, and five frequency domains for the sleep group: delta (0.75-4 Hz), theta (4.25-8), alpha (8.25-13), sigma (13.25-16), and beta (16.25-31 Hz) frequency ranges. In order to reduce the number of parameters, we averaged bandwise spectral power measures of Frontal (frontal: Fp1, Fpz, Fp2, AF3, AF4, F7, F5, F3, F1, Fz, F2, F4, F6, F8, frontocentral and frontotemporal: FT7, FC5, FC3, FC1, FC2, FC4, FC6, FT8), Central (central, centrotemporal and centroparietal: T7, C5, C3, C1, Cz, C2, C4, C6, T8, CP5, CP3, CP1, CPz, CP2, CP4, CP6, TP8), and Posterior (parietal, parietotemporal and occipital: P7, P5, P3, Pz, P2, P4, P6, P8, POz, O1, Oz, O2) electrode derivations.

We quantified sleep spindling activity by the Individual Adjustment Method (IAM, (Bódizs et al., 2009; Ujma et al., 2015) that considers individual spectral peaks to detect spindles in each participant. This method defines frequency boundaries for slow and fast spindles based on the spectral power of NREM sleep. These individualized boundaries are used as frequency limits for slow and fast spindle bandpass filtering (FFT-based, Gaussian filter, 16 s windows) of the EEGs. Thresholding of the envelopes of the band-pass filtered recordings are performed by individual and derivation-specific amplitude criteria (See the description of the method in more detail in Bódizs et al., 2009; Ujma et al., 2015). We used spindle density (spindles/min) and the average amplitude (μV) of slow and fast spindles as different measures of spindling activity. To reduce the number of statistical comparisons, we averaged spindle measures of Frontal, Central, and Posterior electrode derivations similarly to spectral power measures.

### Statistical analyses

Statistical analyses were carried out with the Statistical Package for the Social Sciences version 22.0 (SPSS, IBM) and R (Team, 2014). The blocks of the explicit ASRT task were collapsed into epochs of five blocks to facilitate data processing and to reduce intra-individual variability. The first epoch contained blocks 1-5, the second epoch contained blocks 6-10, etc. We calculated median reaction times (RTs) for all correct responses, separately for pattern, random high and random low trials for each epoch and each participant. Note that for each response (n), we defined whether it was the last element of a high- or a low-frequency triplet. Two kinds of low-frequency triplets were eliminated: repetitions (e.g., 222, 333) and trills (e.g., 212, 343). Repetitions and trills corresponded to low frequency triplets for all participants and individuals often show pre-existing response tendencies to such triplets (Howard et al., 2004). By eliminating these triplets, we attempted to ensure that differences between high vs. low-frequency triplet elements emerged due to learning and not to pre-existing response tendencies.

To show the performance trajectories of RTs for different trial types, and to explore their differences, we performed a mixed design analyses of variance (ANOVA) with EPOCH (1-8) and TRIAL TYPE (pattern, random high, random low) as within-subject factors, and GROUP (Active Wake, Quiet Rest, Nap) as a between-subject factor. To evaluate the effect of epoch and trial type we performed post-hoc comparisons (Fisher’s LSD).

In order to examine the changes in Statistical and Sequence Learning that occur during the training phase, we applied a mixed-design ANOVA with EPOCH (1 −5) and LEARNING TYPE (Statistical Learning, Sequence Learning) as within-subject factors, and GROUP (Active Wake, Quiet Rest and Nap) as a between-subject factor. Post-hoc comparisons were applied to evaluate changes in performance during the training phase in case of Sequence and Statistical Learning.

To examine off-line changes occurring between testing and retesting sessions we used a similar mixed-design ANOVA with EPOCH (6-8) and LEARNING TYPE (Statistical Learning, Sequence Learning) as within-subject factors, and GROUP (Active Wake, Quiet Rest and Nap) as a between-subject factor. Post-hoc comparisons were run to contrast performances of the testing phase (6^th^ epoch) and the retesting phases (7^th^ and 8^th^ epochs).

Greenhouse-Geisser epsilon (ε) correction was used if necessary. Original *df* values and corrected *p* values (if applicable) are reported together with partial eta-squared (η^2^) as a measure of effect size.

Finally, we aimed to examine the associations between EEG spectral power measured during the off-line period and change in learning performance across the testing and retesting phase, in each group separately. Off-line changes in Sequence and Statistical Learning were defined as the difference between the learning scores of the first retesting (7^th^ epoch) session and the testing session (6^th^ epoch). Thus, a positive value indicated improvement in learning performance after the off-line period. Furthermore, we aimed to examine whether EEG spectral power measured during off-line periods predicted additional performance change after longer re-learning, therefore, we calculated a secondary off-line change score contrasting learning scores of the 8^th^ (2^nd^ half of the retesting session) with those of the 6^th^ epoch (testing session).

The associations between sleep spindles and off-line changes of the above measures were also examined (within the sleep group only). Pearson correlation coefficients or (if normality was violated) Spearman rank correlations were run between spectral power values (of each region and band) and off-line changes in learning scores. The issue of multiple comparisons was addressed by the False Discovery Rate correcting for type 1 error (Benjamini and Hochberg, 1995).

## Results

### Group characteristics

Groups were matched in age, gender, working memory, executive function, and initial sleepiness and stress level (Table 1). However, after the one hour long off-line period, the groups differed in sleepiness (*F*_2,75_ = 3.19, *p* = 0.05). Post-hoc test showed that the Nap group scored significantly higher on the KSS (indicating lower sleepiness on the Hungarian version of the KSS scale where higher scores indicate a more refreshed state, that is, lower sleepiness) than the Active Wake group (*p* = 0.02), however the difference was not significant after FDR correction.

Sleep parameters of the Nap group are listed in Table 2. In the Nap group, only one participant reached REM phase during sleep, thus we only report the characteristics of Non-REM sleep.

**Table 2.** Descriptive characteristics of sleep parameters in the Nap group

| Variable | Mean (SD) |
| --- | --- |
| Sleep duration (min) | 41.16 (12.35) |
| Sleep efficiency (%) | 70.28 (16.27) |
| Wake duration (min) | 16.53 (7.77) |
| S1 duration (min) | 6.02 (3.62) |
| S2 duration (min) | 17.93 (6.59) |
| SWS duration (min) | 16.89 (12.82) |
| Fr. fast spindle density | 6.37 (0.96) |
| Cent. fast spindle density | 7.45 (0.83) |
| Post. fast spindle density | 7.35 (0.93) |
| Fr. fast spindle amp. | 4.56 (1.32) |
| Cent. fast spindle amp. | 6.01 (1.56) |
| Post. fast spindle amp. | 5.38 (1.38) |
| Fr. slow spindle density | 7.31 (1.12) |
| Cent. slow spindle density | 7.33 (1.19) |
| Post. slow spindle density | 7.4 (1.16) |
| Fr. slow spindle amp. | 3.91 (1.85) |
| Cent. slow spindle amp. | 3.28 (1.49) |
| Post. slow spindle amp. | 2.54 (0.96) |

*Note* S1 – Stage 1, S2 – Stage 2, SWS – Slow Wave Sleep

### Are performance trajectories of responses to different trial types different between groups?

Overall, participants in the different groups responded with similar RTs (main effect of GROUP: *F*_2,75_ = 0.80, *p* = 0.46, partial *η*^2^ = 0.02). Irrespectively of trial types, RTs significantly decreased across epochs (main effect of EPOCH: *F*_7,525_ = 175.26, *p* < 0.0001, partial *η*^2^ = 0.70), indicating general skill improvements due to practice (Figure 2). The GROUP x EPOCH interaction was not significant (*F*_14,525_ = 1.18 *p* = 0.32, partial *η*^2^ =0.03), suggesting that general skill improvements were similar in the groups. Furthermore, participants showed significant Sequence and Statistical Learning (main effect of TRIAL TYPE: *F*_2,150_ = 52.04, *p* < 0.0001, partial *η*^2^ = 0.41): they responded faster to pattern than random high trials (*p* < 0.0001), and faster to random high compared to random low trials (*p* < 0.0001). The GROUP x TRIAL TYPE interaction was not significant (*F*_4,150_ = 0.80, *p* = 0.46, partial *η*^2^ =0.02) indicating that there was no difference between the groups in performance for different trial types. In addition to that, the EPOCH x TRIAL TYPE interaction was significant (*F*_14,1050_ = 11.93, *p* < 0.0001, partial *η*^2^ = 0.14), indicating different learning trajectories in case of the three trial types (see Figure 2). Although participants became faster for all trial types during the course of the task, responses to pattern trials showed greater gains in comparison to both random trials: Average reaction times of pattern trials decreased from 357.89 to 257.56 ms (*p* < 0.0001), of random high trials from 370.98 to 326.14 ms (*p* < 0.0001), and of random low trials from 388.26 to 349.65 ms (*p* < 0.0001). Practice-dependent improvement in response to pattern trials was significantly higher than the improvement in case of random high (*t*_77_ = 4.81, *p* < 0.0001) and random low (*t*_77_ = 5.45, *p* < 0.0001) trials. The improvement in responses to random high and random low trials was only marginally different (*t*_77_ = 1.84, *p* = 0.07). The GROUP x EPOCH x TRIAL TYPE interaction was not significant (*F*_28,1050_ = 0.66, *p* = 0.68, partial *η*^2^ =0.02), suggesting that performance trajectories to the different trial types were similar among the groups.

**Figure 2.**
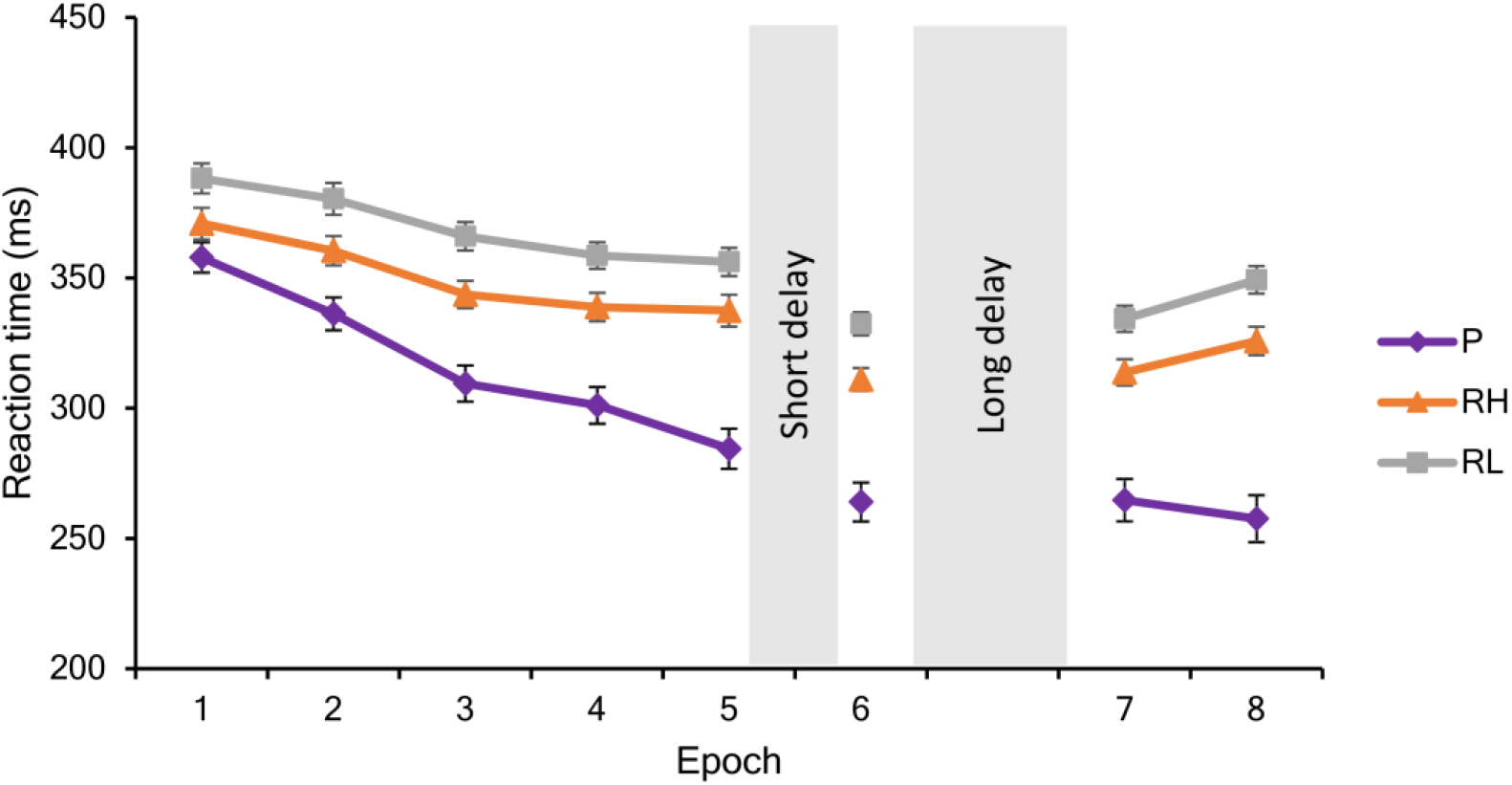
Performance during the training, testing and retesting sessions. Mean reaction times and standard errors are visualized in response to pattern (P), random high (RH), and random low (RH) trials during each epoch.

### Do Sequence and Statistical Learning during training differ between groups?

Sequence and Statistical Learning during the training phase were similar across the groups (main effect of GROUP: *F*_2,75_ = 1.10, *p* = 0.34, partial *η*^2^ = 0.03). Irrespectively of learning type, performance improved across epochs of training (main effect of EPOCH: *F*_4,300_ = 10.92, *p* < 0.0001, partial *η*^2^ = 0.13). The GROUP x EPOCH interaction was not significant (*F*_8_,3_00_ = 0.59, *p* = 0.68, partial *η*^2^ =0.02), suggesting that improvement during training was similar between the groups. In addition, the main effect of LEARNING TYPE was significant (*F*_1,75_ = 3.93, *p* = 0.05, partial *η*^2^ = 0.05): participants showed greater Sequence Learning compared to Statistical Learning (*M* = 32.50 vs. *M* = 19.64, *p* < 0.0001). The GROUP x LEARNING TYPE interaction was not significant (*F*_2,75_ = 0.81, *p* = 0.45, partial *η*^2^ =0.02), suggesting that the difference between Sequence and Statistical Learning were similar among the groups. Furthermore, a significant interaction between EPOCH and LEARNING TYPE emerged (*F*_4,300_ = 5.52, *p* = 0.002, partial *η*^2^ = 0.07): as illustrated in Figure 3, participants, on average, exhibited a steep increase in Sequence Learning during the training phase (the average learning score increased from 13.09 to 53.31 from the 1^st^ epoch to the 5^th^ (*p* < 0.001), whereas Statistical learning occurred in the beginning of the task and remained unchanged by the end of the training phase (the average learning score increased from 17.28 to 18.64 from the 1^st^ epoch to the 5^th^, *p* = 0.68). The GROUP x EPOCH x LEARNING TYPE interaction was not significant (*F*_8,300_ = 0.58, *p* = 0.72, partial *η*^2^ =0.02), suggesting that training-dependent patterns of Sequence Learning and Statistical Learning were similar across the groups.

**Figure 3.**
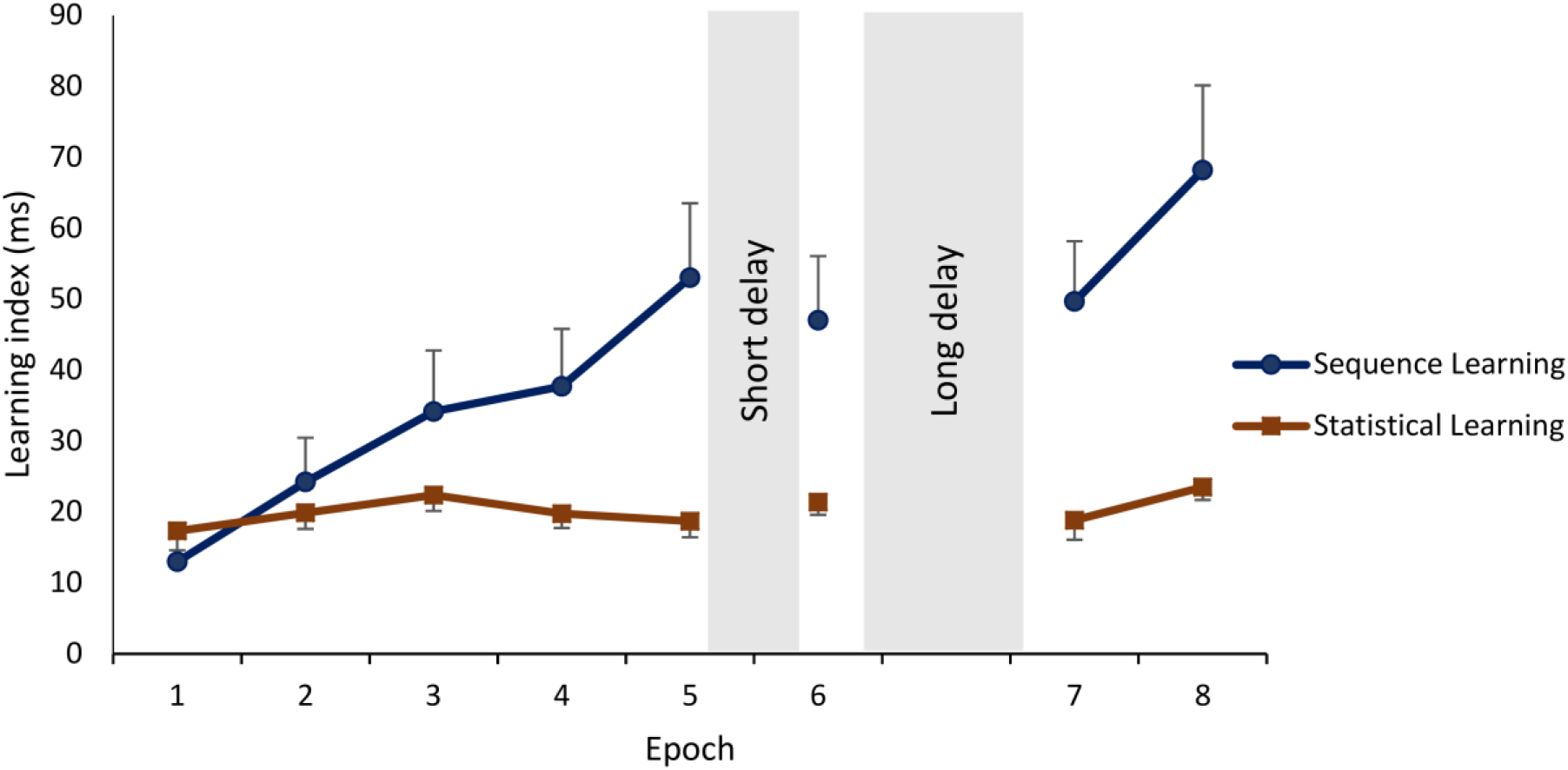
Learning and off-line changes in Sequence and Statistical Learning. Sequence Learning is quantified as the difference in reaction times to random high elements vs. pattern elements. Statistical Learning is quantified the difference in reaction times to random low elements vs. random high elements. Means and standard errors of Sequence Learning and Statistical Learning during each epoch. Sequence Learning exhibited a steep increase during training and additional practice after the off-line periods, whereas Statistical Learning remained unchanged throughout the sessions.

Beyond the group-level results presented in the previous paragraph, we performed an additional analysis to reveal learning trajectories on a subject-by-subject basis. We categorized each subject’s learning trajectory during training by a combination of curve fitting and visual inspection. For comparability, we performed the same steps for Sequence and Statistical learning (see Figure 4A and 4B, respectively) and found that ~33% of participants showed gradually increasing Sequence learning during training, while the trajectory for Statistical learning was gradually increasing only in ~16% of participants (*χ*^2^(1) = 3.80, *p* = 0.05). Compared to these percentages, a relatively smaller number of participants exhibited a step-like increase in learning performance: ~10% of participants for Sequence learning and ~4% of participants of Statistical learning (*p* = 0.15). Additionally, a small portion of participants exhibited a decreasing pattern, with the best performance at the beginning of the task (~5% of participants for Sequence learning, and ~13% of participants for Statistical learning; *p* = 0.42). The learning trajectory of the majority of participants did not clearly follow any of the patterns described above. These learning trajectories were categorized as ‘Other pattern’ (~53% of participants for Sequence learning, and ~66% of participants for Statistical learning; *p* = 0.81). These participants exhibited relatively large changes in performance from one epoch to another and then returned to the previous performance level. The timing of these larger changes in performance was evenly distributed across epochs. It is plausible that these participants explored different (explicit or implicit) strategies over the course of learning that may have resulted in large changes in some epochs compared to their overall learning performance. Note, however, that the primary focus of our study was not to test these possible strategies but to compare Sequence and Statistical learning trajectories across the three experimental groups (Quiet Rest, Active Wake and Nap). Importantly, the distribution of subgroups exhibiting different learning trajectories was similar across the three experimental groups both for Sequence learning (*χ*^2^ (6) = 0.91, *p* = 0.99) and for Statistical learning (*χ*^2^(6) = 1.98, *p* = 0.92).

**Figure 4.**
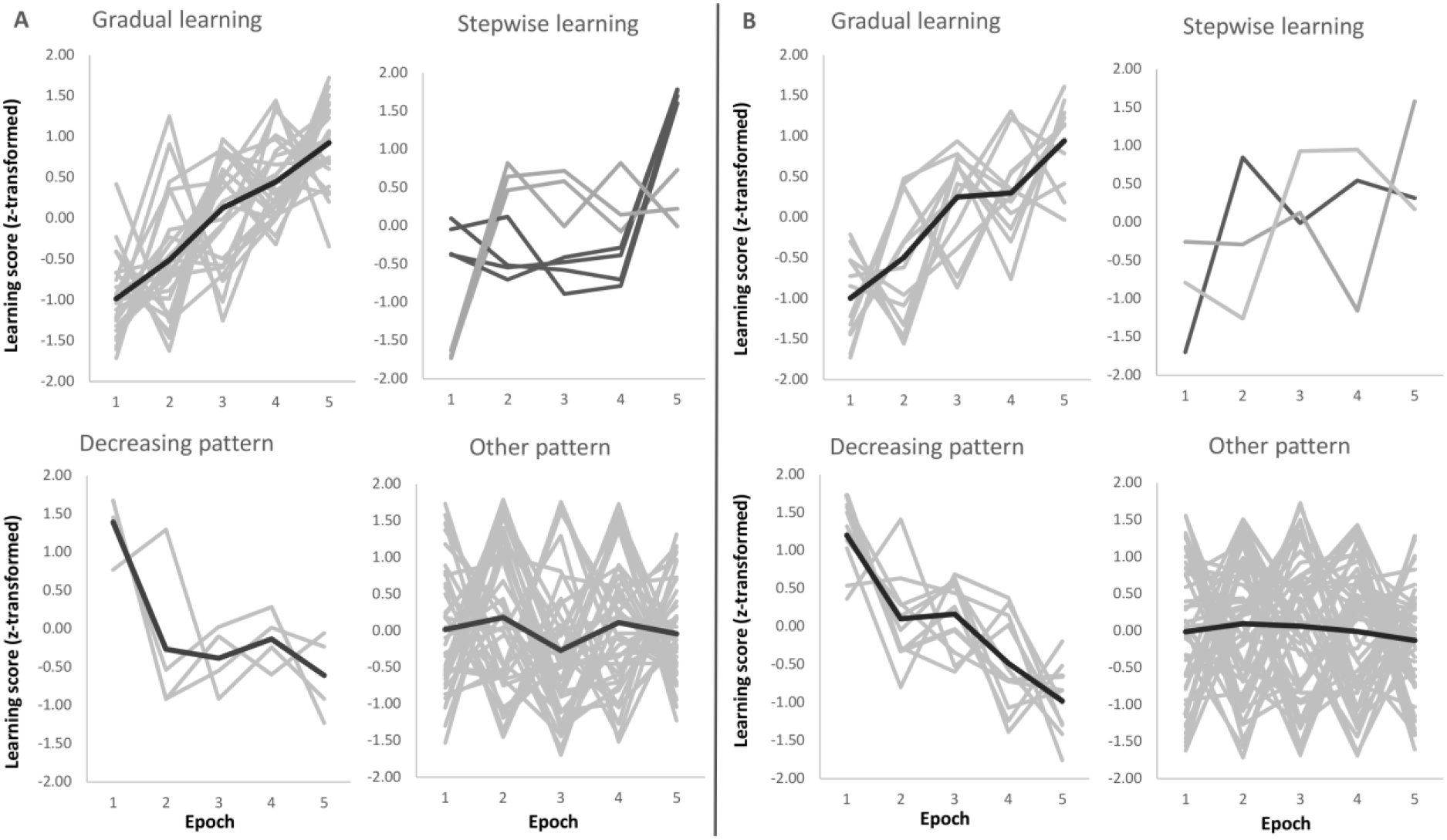
Sequence (A) and Statistical (B) learning trajectories for individual subjects. Each participant’s learning trajectory is presented in a light grey color, while the average learning trajectory for that subgroup is presented in a darker grey color for the ‘Gradual learning’, ‘Decreasing pattern’ and ‘Other pattern’ panels. For the ‘Stepwise learning’ panel, the light and dark grey colors represent subgroups of participants depending on the timing of their performance increase (no average learning trajectory is presented).

### Early Statistical learning effects during training

To provide further insights into the trajectory of Statistical learning, we performed additional analyses by focusing on block-level and below block-level data. The first set of analyses aimed to determine the time point when participants successfully extracted the statistical regularities from the stimulus stream. First, we computed Statistical learning scores for each block of Epoch 1, and tested if these Statistical learning scores were significantly different from zero. We found significant Statistical learning effect already in Block 1 of the ASRT task (*t*(73) = 2.12, *p* = 0.04, Cohen’s *d* = 0.25). Next, we zoomed into Block 1 to further test this learning effect. In this analysis, we split Block 1 into two halves and computed Statistical learning scores for each participant, for each half. This level of granularity seemed the most appropriate so that all participants had at least a few random-high trials (~ 4 trials on average, ranging from 2 to 9), enabling us to compute learning scores for all participants. These Statistical learning scores were submitted into one sample t-tests, which showed that Statistical learning scores did not reach significance in the first half of Block 1 (*t*(73) = 1.11, *p* = 0.269, Cohen’s *d* = 0.13), while they were significant in the second half of Block 1 (*t*(73) = 1.99, *p* = 0.05, Cohen’s *d* = 0.23). This analysis thus demonstrates that statistical regularities are learned (albeit very quickly) and the observed significant Statistical learning scores at the very early phase of the task are not due to other (not learning-related) preexisting tendencies.

This rapid learning effect is in fact not surprising if we consider that 80 trials are presented in the first block, and ~ 50 of those trials can be categorized as high frequency triplets (occurring in pattern or random positions). As there are 16 individual triplets that are high frequency, that means that participants encounter each individual triplet approximately four times in the first block already. In contrast, there are 48 individual triplets that are low frequency, and participants encounter these individual triplets approximately (or less than) once in a block. Thus, the observed significant Statistical learning scores (i.e., the difference between the random-high and random-low frequency trials) suggests that participants are so sensitive to the frequency statistics that as little as, on average, four presentations of the same trials are sufficient to show speeded responses to them.

Nevertheless, it is important to highlight that significant learning does not necessarily mean that participants have a stable knowledge about the statistical regularities. Thus, even though the Statistical learning scores are already significant at the early phase of learning and these scores numerically do not change as the task progresses, it is reasonable to assume that more practice can help strengthen the acquired knowledge. We ran an additional analysis to test this assumption. In this analysis, we focused on block-level data and computed Cohen’s *d* effect sizes for the block-level Statistical learning scores. These effect sizes were substantially smaller in the first five blocks of the ASRT task (0.27 on average for Blocks 1-5, i.e., Epoch 1) compared to the later blocks (blocks of Epoch 2: 0.45, Epoch 3: 0.51, Epoch 4: 0.53, Epoch 5: 0.50). This difference in the effect sizes suggests that, although participants were able to extract the statistical regularities from the stimulus stream very early in the task, additional training helped them strengthen the acquired statistical knowledge.

### Are off-line changes in Sequence and Statistical Learning different across the groups?

The three groups did not show different patterns of Sequence and Statistical Learning from the testing to the retesting sessions, as neither the main effect of GROUP (*F*_2,75_ = 0.65, *p* = 0.53, partial *η*^2^ = 0.02), nor the interactions GROUP x EPOCH (*F*_4150_ = 0.52, *p* = 0.67, partial *η*^2^ = 0.01), GROUP x LEARNING TYPE (*F*_2,75_ = 0.65, *p* = 0.53, partial *η*^2^ = 0.02), and GROUP x EPOCH x LEARNING TYPE (*F*_4,150_ = 0.73, *p* = 0.55, partial *η*^2^ = 0.02) emerged as significant predictors. The lack of a group effect is shown in Figure 5 that illustrates offline changes (7^th^ minus the 6^th^ epoch) in Sequence and Statistical Learning separately for each group. Similarly to the training phase, participants exhibited higher scores in Sequence Learning than in Statistical Learning (main effect of LEARNING TYPE: *F*_175_ = 10.72, *p* = 0.002, partial *η*^2^ = 0.13). Moreover, learning indices produced robust changes across epochs as indicated by a significant main effect EPOCH (*F*_2,150_ = 18.99,*p* < 0.0001, partial *η*^2^ = 0.20). More specifically, overall performances (regardless of learning type) were unchanged from the testing phase (6^th^ epoch) to the first retesting epoch (7^th^) (*p* = 0.86), but improved (*p* < 0.0001) from the testing phase to the end of the retesting session (8^th^ epoch), and from the first retesting epoch to the second (7^th^ epoch vs 8^th^ epoch) (*p* < 0.0001). Furthermore, Sequence Learning and Statistical Learning scores showed different patterns after the off-line period (see Epoch 7 and 8 in Figure 3), as indicated by the significant EPOCH x LEARNING TYPE interaction (*F*_2,150_ = 5.31, *p* = 0.009, partial *η*^2^ = 0.07). Neither Sequence Learning nor Statistical Learning seemed to show immediate (early) gains after the off-line period. Sequence Learning scores did not significantly change from the testing phase to the first epoch of retesting (6^th^ epoch, *M* = 47.02 vs. 7^th^ epoch, *M* = 47.69, *p* = 0.85). Similarly, Statistical Learning remained unchanged from testing to the first retesting (6^th^ epoch, *M* = 21.39 vs. 7^th^ epoch, *M* = 19.96, *p* = 0.56). Nevertheless, additional practice produced robust changes in Sequence Learning, that increased significantly from the testing phase to the second epoch of the retesting phase (8^th^ epoch, M= 68.19, *p* = 0.001), whereas Statistical Learning did not show any significant changes by the end of the retesting phase (8^th^ epoch: *M* = 23.51, *p* = 0.41).

**Figure 5.**
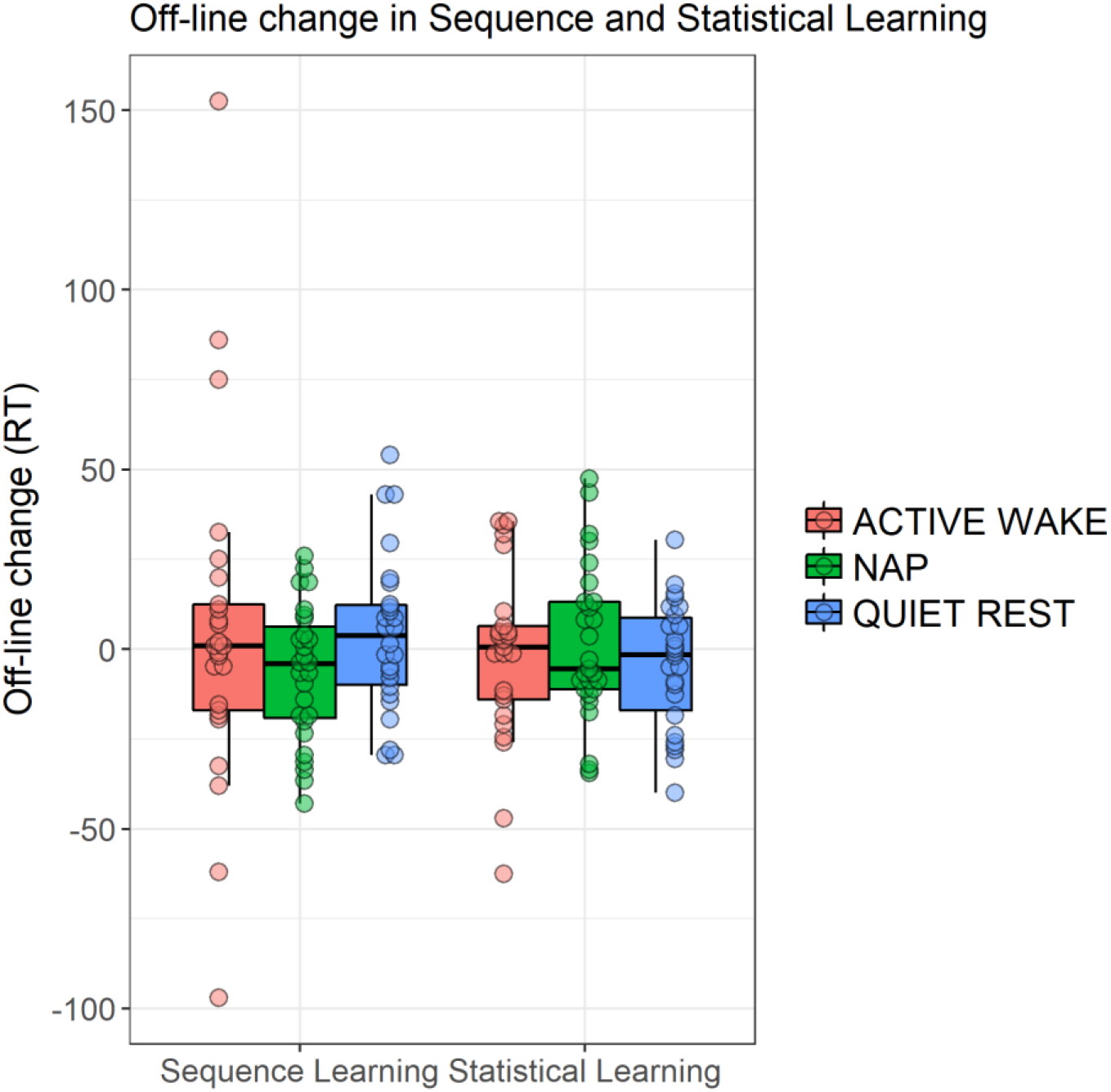
Off-line changes in learning indices within the three groups. Off-line changes were calculated by the learning scores of the 7^th^ epoch minus the respective learning scores of the 6^th^ epoch. Dots show individual data points, the vertical line within the boxes show the medians, boxes represent the first and third quartiles, whiskers indicate the interquartile range of 1.5.

To further explore potential group differences during the off-line period we ran additional ANOVAs separately for Sequence and Statistical learning scores considering their different learning curves. Based on these ANOVAs, we found no group differences in the consolidation (6^th^ epoch vs. 7^th^ epoch) of the acquired knowledge (Sequence learning: *p* = 0.35, Statistical learning: *p* = 0.78). Similarly, no group differences emerged in the additional increase between 7^th^ epoch and 8^th^ epoch (Sequence learning: *p* = 0.65, Statistical learning: *p* = 0.36).

### Awareness of the sequence in the groups

For the analysis of sequence awareness, two participants’ data had to be excluded due the technical issues during collection of sequence reports (one data from the active wake and one data from the nap group). Additionally, eleven participants could not report the correct sequence consistently during training (*N* = 3 in the active wake, *N* = 3 in the nap, and *N* = 5 in the quiet rest group), and therefore they were also excluded from the following analyses. Importantly, there were no group differences in the number of participants who could or could not report the correct sequence consistently and were excluded (chi-square = 1.77, *p* = 0.78).

On average, participants could report the correct sequence consistently from the 6^th^ block (*M* = 6.58, *SD* = 7.04), with no differences across the groups (*F*_2,64_ = 1.53, *p* = 0.23). Overall, the block number from which participants could consistently report the correct sequence showed a significant negative correlation with the Sequence learning scores (*r* = − 0.28, *p* = 0.02). Thus, the earlier participants could find the correct sequence and report consistently thereafter, the better their overall Sequence learning was. No association was observed between the block number and the Statistical learning scores (*r* = −0.06, *p* = 0.63), suggesting that sequence awareness primarily affected Sequence learning but not Statistical learning.

Finally, we conducted an ANOVA for the Sequence learning scores of the training phase (Epoch 1 to 5), including the block number from which participants could consistently report the correct sequence as a covariate to check how sequence awareness affected the time course of learning across groups. The ANOVA revealed a significant main effect of EPOCH (*F*_4,244_ = 10.53, *p* < 0.001, partial *η*^2^ = 0.147), indicating better Sequence learning scores as learning progressed. This effect was modulated by the block number on a trend level (*F*_4,244_ = 2.58, *p* = 0.08, partial *η*^2^ = 0.041), suggesting that the earlier participants could report the correct sequence, the better their Sequence learning became across training. Importantly, no significant group differences emerged either in overall learning or in the trajectory of learning even after taking into account the block number as a covariate (*p*s > 0.21).

A similar ANOVA was conducted for the consolidation analysis (Epoch 6 to 8). This ANOVA also revealed a significant main effect of EPOCH (*F*_2,122_ = 8.34*,p* < 0.001, partial *η*^2^ = 0.120), which is consistent with the previous ANOVA conducted for these epochs, showing increase in Sequence learning scores due to additional training (Epoch 7 vs. Epoch 8, see Figure 3). This effect was not modulated by the block number (*p* = 0.49). Furthermore, no significant group differences emerged either in overall learning scores or in the trajectory of learning scores across these epochs, even after taking into account the block number as a covariate (*p*s > 0.32). These results altogether suggest that, although the timing when participants gained explicit knowledge about sequence affects their Sequence learning scores, this effect is similar across the groups both during training and consolidation.

### Associations between EEG spectra and off-line changes

Off-line changes in Sequence Learning as indexed by the difference scores between the 7^th^ (first half of retesting phase) and the 6^th^ epochs’ (testing phase) scores were positively associated with frontal theta power (*r* = 0.44 *p* = 0.028) within the nap group. Off-line changes in Sequence Learning were not associated with spectral EEG power measures in the either of the awake (AW, QR) groups. Additional off-line-changes in Sequence Learning as indexed by the difference scores between the 8^th^ (second half of retesting phase) and the 6^th^ epochs’ (testing phase), showed a positive association with frontal theta power (*r* = 0.52, p= 0.008) within the nap group only. Nevertheless, these correlations did not reach statistical significance after FDR correction of multiple comparisons (all *p*s > 0.05). Since region-wise averaging of electrodes might not capture associations between behavioral measures and spectral power of a more local nature, we examined (on an exploratory level) the associations between theta activity and off-line changes (7^th^ vs 6^th^ epoch and 8^th^ vs 6^th^ epoch) in Sequence Learning within the nap group. As shown in Figure 6, associations with theta band power were prominent at frontal electrode sites, peaking at left frontopolar locations in case of immediate off-line changes (Figure 6A), as well as in case of additional off-line changes in performance (Figure 6B). Finally, we examined the associations between off-line (7^th^ vs 6^th^ epoch and 8^th^ vs 6^th^ epoch) changes in Sequence Learning and bin-wise EEG spectral power averaged across all electrodes (within the Nap group). Immediate (7^th^ vs 6^th^ epoch) and delayed (8^th^ vs 6^th^ epoch) post-sleep improvement in Sequence Learning correlated only with slow frequency activity between 2-7.75 Hz (all bins *p* < 0.01).

**Figure 6.**
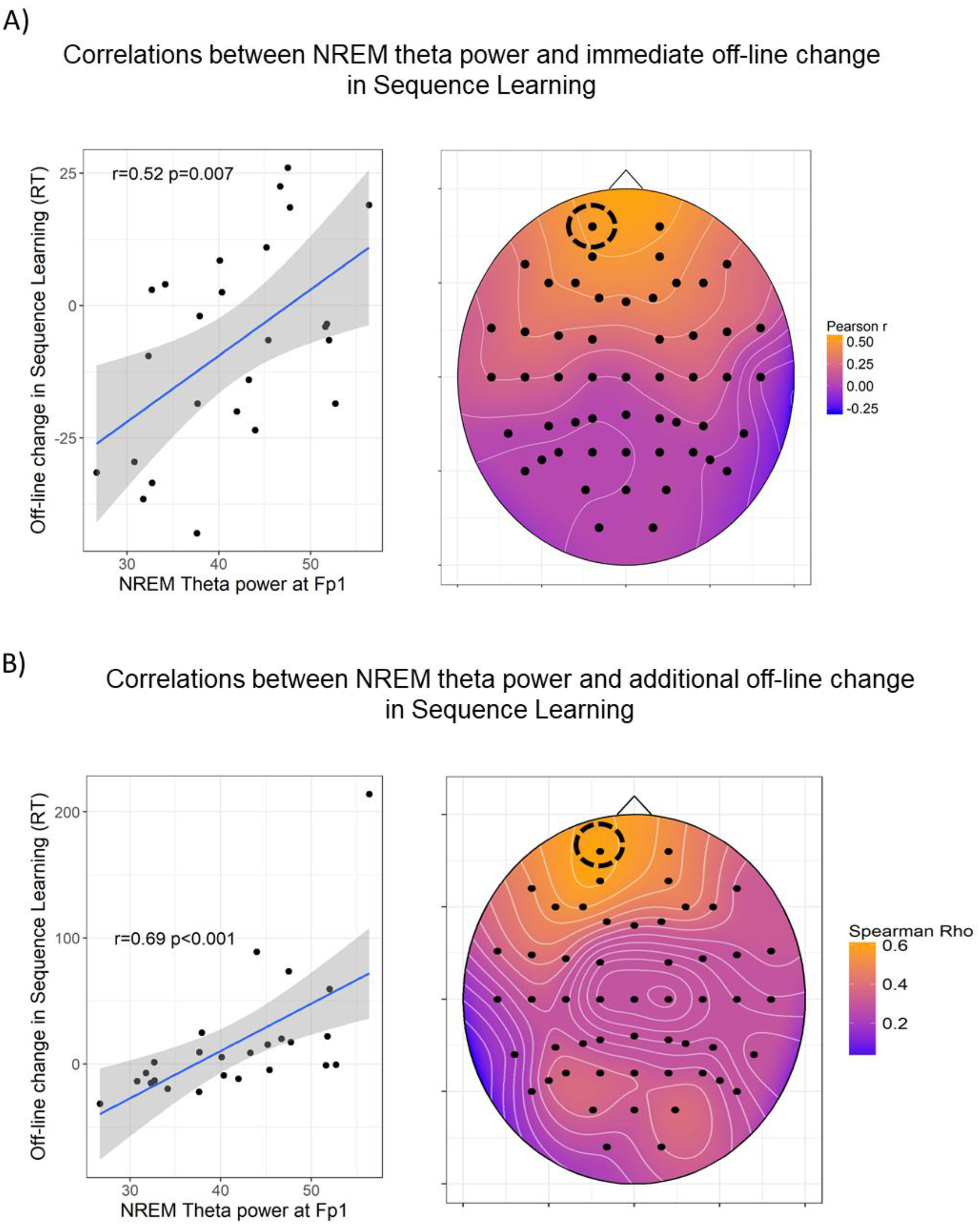
Associations between NREM theta power and off-line changes in Sequence Learning. A) Pearson correlations between NREM theta band power and immediate (7^th^ vs 6^th^ epoch) post-sleep changes in Sequence Learning. B) Spearman Rho correlations coefficients between NREM theta band power and delayed (8^th^ vs 6^th^ epoch) post-sleep changes in Sequence Learning. The heat plots on the right indicate the magnitude of correlation coefficients, the scatterplots on the left show the association in a prominent (left frontal) electrode site. In case of 5B the correlation coefficient remained unchanged (*r* = 0.64, *p* < 0.001) after the exclusion of the outlier. The figures show uncorrected *p* values (before FDR correction). For the immediate off-line changes, only Fp1, Fp2, AF3, AF4 locations remained significant after FDR correction. For the additional off-line changes, frontal channels Fp1, Fpz, Fp2, AF3, AF4, F7, F5, F3, F1, Fz, F2, F4, F6, F8 as well as FC4, FC5, CP5 and P5 locations remained significant after FDR correction.

Immediate and additional off-line changes (7^th^ vs 6^th^ epoch and 8^th^ vs 6^th^ epoch) in Statistical Learning were not associated with spectral power measures within the nap group, and no other associations emerged within the Quiet Rest and Active Wake groups.

In sum, individual differences in off-line changes in Statistical Learning assessed immediately after the long delay (6^th^ vs. 7^th^ epoch) and after extended practice, (6^th^ vs. 8^th^ epoch) were not associated with spectral EEG power measures in any of the three groups. On the other hand, immediate and delayed post-sleep improvements in Sequence Learning were predicted by high delta and theta activity during sleep within the Nap group. Nevertheless, these correlations did not remain significant after correction for multiple comparisons.

### Associations between sleep spindles and off-line changes

Off-line change (7^th^ vs 6^th^ epoch) in Sequence Learning showed a negative correlation with slow spindle density at Frontal (*r* = −0.52, *p* = 0.008), Central (*r* = −0.54, *p* = 0.006) and Posterior (*r* = −0.53, *p* = 0.006) derivations. Slow spindle amplitude, fast spindle density and amplitude were not associated with the off-line change in Sequence Learning. Negative correlations between slow spindle density and off-line change in Sequence Learning remained significant after FDR correction (*p* = 0.036).

Off-line change in Statistical Learning was negatively correlated with fast spindle amplitude (Frontal: *r* = −0.43, *p* = 0.03; Central: *r* = −0.47, *p* = 0.02; Posterior: *r* = −0.44, *p* = 0.03), but was not related either to fast spindle density or slow spindle density/amplitude. Correlations between fast spindle amplitude and off-line change in Statistical Learning were not significant after FDR correction (all *p*s > 0.05).

To examine whether the negative correlation between off-line changes in performance and spindle parameters were linked to overall Sequence/Statistical Learning ability, we applied partial correlations with learning performance of the training phase as a covariate. Learning performance here was computed as the differences in Sequence and Statistical learning between the 5^th^ and the 1^th^ epochs of the training phase. Slow spindle density remained a negative correlate of off-line change in Sequence Learning even after controlling for this initial Sequence Learning performance (Frontal: *r* = −0.5, *p* = 0.006; Central: *r* = − 0.52, *p* = 0.009; Posterior: *r* = −0.51, *p* = 0.005).

Similarly, partial correlations were computed between fast spindle amplitude and offline change in Statistical Learning with Statistical Learning performance as a covariate. The correlations showed trends after partialing out this initial Statistical Learning performance (Frontal: *r* = −0.37, *p* = 0.07; Central: *r* = −0.43, *p* = 0.03; Posterior: *r* = −0.36, *p* = 0.08).

Additional (delayed) off-line-changes in Sequence and Statistical Learning as indexed by the difference scores between the 8^th^ (second half of retesting phase) and the 6^th^ epochs’ (testing phase) were not associated to any of the extracted spindle parameters.

## Discussion

Our aim was to investigate performance trajectories in Sequence and Statistical Learning during extensive practice and after off-line periods spent in different vigilance states. In order to examine these processes in the same experimental context, we applied a paradigm that simultaneously measured sequence and statistical learning by delineating order and frequency-based information. Our findings indicate that Sequence and Statistical Learning follow different learning curves. Whereas performance in Sequence Learning exhibited an increase during training, Statistical Learning was rapidly acquired and remained unchanged throughout training. During the off-line period, both forms of learning were preserved as no significant off-line changes emerged in either Sequence or Statistical Learning. Nevertheless, Sequence Learning improved after additional practice (i.e., in the retesting phase), whereas Statistical Learning remained stable regardless of further training compared to the testing phase. Performance trajectories were similar across the groups: Performance during training and consolidation did not differ between the Active Wake, Quiet Rest and Nap groups. EEG spectral power assessed during the off-line periods was not associated with off-line changes in Sequence and Statistical Learning in the awake groups. Within the Nap group we found a trend indicating a positive association between frontal theta band power and off-line change in Sequence Learning. In addition, frontal theta power predicted further improvements in Sequence Learning after additional practice. Within the Nap group, slow spindle density was negatively associated with post-sleep improvement in Sequence Learning, and fast spindle amplitude was negatively associated with post-sleep improvement in Statistical Learning.

Our data suggests that sequence and statistical learning are markedly different subprocesses of procedural learning. Frequency-based information is acquired rapidly and appears to undergo less prominent changes during further training compared to the acquisition of order-based information that may exhibit further performance improvements. Our fine-grained analyses revealed that statistical learning occurs already in the first block of the task. This finding suggests that participants are so sensitive to the frequency statistics that as little as, on average, four presentations of the same trials are sufficient to show speeded responses to them. Nevertheless, the further analysis of effect sizes showed that, although participants were able to extract the statistical regularities from the stimulus stream very early in the task, additional training helped them strengthen the acquired statistical knowledge.

Rapid statistical learning has also been reported before: for instance, in the ASRT study of Szegedi-Hallgató and colleagues (2017), statistical learning was apparent already in the first epoch in the Explicit group but seemed to have larger individual differences in the Implicit groups as only one of the two Implicit groups exhibited significant statistical learning in the first epoch (see Supplementary results and figures). Similarly, in Kóbor et al.’s (2018) study, statistical learning was observed in the first epoch of the explicit version of the ASRT task, along with a significant sequence learning as well. Consequently, a possible explanation for the very rapid statistical learning is that, in an explicit condition, the instructions and motivation to learn can have an overarching effect, providing a cognitive state, in which not only the instructed sequential but also the uninstructed statistical regularities can be learned quickly. Although this was not in the primary focus of these previous studies, if we take a closer look at the learning trajectories, it appears that statistical regularities are extracted very early and no (or very little) further gains may be observed during training if explicit instructions are given for the *sequential* information (Szegedi-Hallgató et al., 2017; Kóbor et al., 2018). In contrast, in the implicit conditions, statistical learning may undergo further improvements during training (Szegedi-Hallgató et al., 2017), above and beyond the strengthening of the acquired knowledge as suggested in the previous paragraph. These observations support the interpretation that explicit instructions and the motivation to learn can have an overarching effect in that not only the instructed sequential but also the uninstructed statistical regularities can be learned more quickly. Interestingly, a recent study showed that, if the task is fix-paced instead of self-paced, no such overarching effect can be observed, suggesting a complex interplay of multiple factors that may influence the effect of explicit instructions on learning (Horvath et al., 2018). Further studies should directly test these factors.

Nevertheless, it is important to note that statistical learning typically occurs implicitly (i.e., without conscious intent to learn and without awareness about the learning situation itself or about the actual regularities) and relatively quickly, already in one learning session (e.g., Song et al., 2007a; Nemeth et al., 2013; Kóbor et al., 2017). In contrast, it has been previously shown that acquiring the alternating sequence structure (frequently referred to as higher-order sequence learning) in the ASRT task typically occurs after 4 days of practice if learning is implicit (Howard and Howard, 1997; Howard et al., 2004), while this can be substantially faster if explicit instruction is provided to the participants (Nemeth et al., 2013). Accordingly, participants quickly formed explicit knowledge about the sequence. Therefore, we think that the current study design was suitable to measure both sequence and statistical learning, bringing them in the same time frame of acquisition (i.e., showing significant learning in one learning session for both measures).

The present study narrows down the concept of statistical learning by regarding it as only one of the processes that is the sensitivity to frequency information. From a theoretical perspective, however, it is important to note that at the level of transitional probabilities, statistical learning (in this narrow sense) and sequence learning could be considered as similar. Namely, both are statistical learning in a broader sense. When acquiring frequency information (statistical learning in the narrow sense), a 2nd order probabilistic sequence should be learned, in which there are always one probable continuation and some less probable continuations for the first two elements of a given three-element stimulus chunk (Szegedi-Hallgató et al., 2017; Kóbor et al., 2018). When acquiring order information (sequence learning), the 2nd order transitional probability is equal to one; namely, consecutive elements in the sequence could be predicted with 100% certainty from the previous sequence element (Kóbor et al., 2018).

Our finding of different learning trajectories within one learning session is in line with the results of Kóbor and colleagues (2018) well as corroborates earlier data (Nemeth et al., 2013) that showed different developmental trajectories of sequence and statistical learning between 11 and 40 years of age but did not analyze the time course of these learning types. Beyond the group-level results, we performed an additional analysis to characterize learning trajectories on a subject-by-subject basis. This analysis revealed that one-third of participants showed gradually increasing Sequence learning during training, and this proportion was significantly higher than the number of participants who exhibited gradually increasing Statistical learning, confirming differences in learning trajectories for Sequence vs. Statistical learning beyond the group-level findings. Nevertheless, the majority of participants exhibited a learning trajectory other than gradual. It is plausible that these participants explored different strategies over the course of learning that may have resulted in large changes in some epochs compared to their overall learning performance. Further investigations should directly focus on individual level heterogeneity and test which factors/characteristics predict learning trajectories on the individual level.

We had a special focus on the off-line change and the effect of sleep on Sequence Learning and Statistical Learning. In order to differentiate between the specific effects of sleep and from the indirect effect of reduced interference during off-line periods, we included a quiet rest control group into the design. On the behavioral level, we found no sleep-dependent consolidation neither in Sequence Learning nor in Statistical Learning. The lack of evidence for the beneficial influence of sleep on statistical learning is in line with previous studies that used probabilistic sequence learning tasks (Peigneux et al., 2003, 2006; Song et al., 2007a; Nemeth et al., 2010; Hallgató et al., 2013), however, we should note that these studies did not differentiate between order-based and frequency-based learning mechanisms. Here, we aimed to investigate the influence of sleep on pure (frequency-based) statistical learning in the perceptual-motor domain. Other studies examined sleep-dependent consolidation on statistical learning in the auditory domain (Durrant et al., 2011, 2013) and contrary to our results, found improved performance after sleep compared to wakefulness. Discrepancies between these studies and our findings might stem from methodological differences (overnight sleep and longer daytime naps in Durrant and colleagues’ study) as well as the examined modality (auditory system vs. perceptual-motor system). Nevertheless, it is important to highlight that Durrant and colleagues (2011) did not include a quiet rest condition that might be favorable in napping studies.

Interestingly, and contrary to our expectations sleep did not facilitate off-line improvement in Sequence Learning either. In case of perceptual-motor sequence learning, Robertson and colleagues (Robertson et al., 2004) reported sleep-dependent consolidation in the explicit version of the Serial Reaction Time task using deterministic sequences. Discrepant findings between the present and Robertson and colleagues’ study can be the result of different sequence structures applied in the SRT and ASRT task. In addition, other confounding factors, such as the effects of fatigue or reactive inhibition (Török et al., 2017) might have a different impact on these tasks. For instance, effects of fatigue are typical to occur in learning tasks (Rickard et al., 2008; Brawn et al., 2010; Pan and Rickard, 2015), however, ASRT learning scores seem to be relatively immune against the influence of fatigue (Török et al., 2017). Furthermore, recent studies raised concerns about the reliability of the deterministic SRT task (Stark-Inbar et al., 2017; West et al., 2017) while the ASRT proved to be a more reliable measure of sequence learning (Stark-Inbar et al., 2017).

Performance in Sequence and Statistical Learning did not show off-line improvements immediately after the long delay period; however, performance in Sequence Learning exhibited further gains after additional practice, suggesting that post-sleep increases in our case were also largely dependent on further practice. Interestingly, delayed (training-dependent) off-line improvements were associated with slow oscillatory activity within the Nap group. This finding suggests that not sleep *per se*, but low-frequency oscillations are associated with delayed performance gains after sleep and additional practice. Our findings indicate that slower oscillatory activity including the (high) delta and the theta frequency ranges (from 2 to 7.75 Hz) during daytime sleep might be predictive of post-sleep improvements in Sequence Learning. Slow frequency oscillations peaking at anterior locations and spanning between 1 Hz to 8 Hz reflect the homeostatic and restorative capacity of sleep as power in these frequencies is increased after prolonged wakefulness (Borbély et al., 1981; Marzano et al., 2010) in fronto-central derivations. Furthermore, the homeostatic increase in spectral power between 2-7 Hz is state-independent (Marzano et al., 2010) making these oscillations likely candidates to reflect restorative processes during a daytime nap, with lower homeostatic pressure. Whether the association between slow frequency activity and further improvement in Sequence Learning reflects processes of sleep-related memory consolidation or a non-specific effect of restorative sleep facilitating performance remains a question of further research.

Sleep spindle parameters within the Nap group were negatively associated with offline changes in performance: slow spindle density and fast spindle amplitude showed negative associations with early off-line changes in Sequence Learning and Statistical Learning, respectively. These findings are hard to interpret as they are at odds with the majority of previous findings that reported a positive association between spindle parameters, general cognitive abilities, and off-line gains in performance in a variety of declarative and procedural learning tasks (see Rasch and Born, 2013 for a comprehensive review). Still, negative correlations were also reported to some extent although in samples including children (Chatburn et al., 2013), and psychiatric patients (Nishida et al., 2016). In our study, associations between spindle parameters and off-line changes in performance might not simply stem from trait-like effects, as associations were unchanged if we controlled for the confounding effects of training-dependent learning performance. Nevertheless, given the lack of baseline EEG measurements, we cannot fully discern trait- and state-like effects in the present study. Moreover, only the association between slow spindle density and the off-line change in Sequence Learning remained significant after the correction for multiple comparisons, whereas previous studies mainly linked sleep-dependent cognitive benefits to fast spindle activity. In sum, off-line changes in Sequence Learning and Statistical Learning were associated with different spindle parameters, nevertheless, the relevance of these associations should be examined in further studies, including baseline sleep measurements without pre-sleep learning experience.

To conclude, here we were able to assess the time-course of two fundamental learning processes, namely Sequence Learning and Statistical Learning separately and showed that Statistical Learning is acquired rapidly and remains unchanged even after extended practice, whereas Sequence Learning may develop more gradually. On the behavioral level, both sequence and statistical knowledge were retained and were independent of whether the offline period included sleep or not. Although our measures of cortical oscillations assessed during the off-line period showed associations with behavioral performance within the sleep group to some extent, the influence of sleep-specific oscillations on Sequence and Statistical learning should be examined in future studies. Nevertheless, our findings suggest that sleep does not have an all-in-one-effect on memory consolidation, and future studies should focus on mapping systematically which learning and memory mechanisms might and might not benefit from sleep and related oscillatory activity. Learning and memory should be assessed on a process level (such as Sequence Learning and Statistical Learning in the current study) in order to characterize the time-course of these processes on the behavioral level as well as their neural correlates more precisely.

## Acknowledgements

This research was supported by the Research and Technology Innovation Fund, Hungarian Brain Research Program (2017-1.2.1-NKP-2017-00002 PI: D.N.); Hungarian Scientific Research Fund (NKFI K 128016, NKFI PD 124148, PI: K.J., NKFI PD 115432, PI: P.S.); and János Bolyai Research Fellowship of the Hungarian Academy of Sciences (to K. J. and P.S.). The authors declare no competing financial interests. This manuscript has been released as a pre-print at BioRxiv (https://www.biorxiv.org/content/early/2018/02/23/216374) (Simor et al. 2017).

Author contributions
DN, KJ and PS conceived the original idea and designed the study. KJ programmed the experimental tasks. ZZ, NE, KH, OP and CT performed the experiments, collected andpreprocessed the data. PS, ZZ, KJ, FG and NÉ analyzed the data. PS, ZZ, KJ and DN wrote the manuscript. All authors discussed the results and commented on the manuscript.

